# An Investigation of Neural Mechanisms Underlying Verb Morphology Deficits in Aphasia

**DOI:** 10.1101/2021.05.28.445987

**Authors:** Madeline Pifer, Christian Brodbeck, Yasmeen Faroqi-Shah

## Abstract

Agrammatic aphasia is an acquired language disorder characterized by slow, non-fluent speech that include primarily content words. It is well-documented that people with agrammatism (PWA) have difficulty with production of verbs and verb morphology, but it is unknown whether these deficits occur at the single word-level, or are the result of a sentence-level impairment. The first aim of this paper is to determine the linguistic level that verb morphology impairments exist at by using magnetoencephalography (MEG) scanning to analyze neural response to two language tasks (one word-level, and one sentence-level). It has also been demonstrated that PWA benefit from a morphosemantic intervention for verb morphology deficits, but it is unknown if this therapy induces neuroplastic changes in the brain. The second aim of this paper is to determine whether or not neuroplastic changes occur after treatment, and explore the neural mechanisms by which this improvement occurs.

## Introduction

> *“Yeah…uh I have-ired a stroke in one-thousand-nine-hundred-ninety-nine….and um reason um. I uhh know six years ago I not know anything um speech… arm or leg umm…. and I need to um work on it…on me” -*AP10, conversational speech sample

Agrammatic aphasia is an acquired language disorder characterized by slow, non-fluent speech with short phrases/sentences that include primarily content words (Menn, Obler, & Miceli, 1990). It is well-documented across a variety of sources that persons with agrammatic aphasia have particular difficulty with production of verbs and verb morphology. While verb naming deficits are a symptom seen broadly in many types of aphasia (including nonfluent) (Matzig, Druks, Masterson, & Vigliocco, 2009), persons with agrammatism seem to be uniquely challenged by verb production (Zingeser & Berndt, 1990), and more specifically, they tend to show difficulty with production of verb inflections, particularly when these are used for marking tense (Faroqi Shah & Thompson, 2007; Friedmann & Grodzinsky, 1997). Deficits with comprehension of verb inflections have also been documented in agrammatism, and have been shown to be correlated with verb tense production deficits (Dickey, Milman, & Thompson, 2005; Faroqi-Shah & Dickey, 2009).

While deficits in production and comprehension of tense morphology at the sentence level have been clearly identified in persons with agrammatic aphasia, it is not known how they process verb inflections in isolation, or if any sentence processing deficits are actually the downstream effect of difficulty with processing inflected verbs. This paper has two aims. The first is to better characterize the impairments in verb inflection processing seen in agrammatic aphasia. Specifically, this study examines the processing of inflected verbs both in isolation and in sentence contexts, by using magnetoencephalography (or MEG) to examine neural activation while persons with agrammatic aphasia and neurotypical adults complete two verb processing tasks. The second aim of this project is to further understand the neural mechanism(s) by which agrammatic persons improve in their verb production following a morphosemantic intervention by analyzing MEG data collected during verb processing tasks.

### Agrammatism and Theories of Verb Deficits

Early predominant views on agrammatism purported that agrammatism is a reflection of a selective syntactic deficit (Berndt & Caramazza, 1980) that is independent of semantic/lexical deficits (Caramazza & Berndt, 1978). A variety of early theories addressed syntactic comprehension deficits in agrammatism, such as the Trace Deletion Hypothesis (Grodzinsky, 1986; Grodzinsky, 1995), the Double Dependency Hypothesis (Mauner, Fromkin, & Cornell, 1993), and theories surrounding impairments in phonological working memory (Linebarger et al., 1983). These theories do not explain the deficits seen in production of verb morphology in agrammatism.

Other theories directly tackle the issue of verb inflection production in agrammatism. Studies of verb inflection in agrammatism have approached the issue by considering either the morphophonological properties of the verb, or the functions served by a particular verb inflection.

The first perspective identifies the morphophonological properties of inflected verbs that serve to increase demands on language production, including phonological complexity and affixation (Obler, Harris, Meth, Centeno, & Mathews, 1999). It has been proposed that inflected verbs constitute complex, compositional words that are subject to a unique morphophonological processing that occurs in the left inferior frontal gyrus and basal ganglia, resulting in problems with verb inflection when this region is damaged (Ullman et al., 1997; 2005). This complex morphophonological processing is unique to verbs and unrelated to general phonological processing (Tyler et al., 2004; Tyler, Randall, & Marslen-Wilson, 2002). Phonological complexity has been investigated by comparing regular and irregular verbs, as well as within irregular verbs, given that irregular verbs fall into different sub-regular families. For instance, in English, regular past tense verbs often end in consonant clusters (e.g., *fixed* and *pushed*), while irregular past tenses may not (*sat* and *dug*, but see *slept* and *told*), making the former more phonologically complex. One study cited greater impairment in sentence completion task for irregular verbs that required a stem change (e.g. sell > sold) (Marusch, Jäger, Burchert, & Nickels, 2017).

When considering verb inflections, some authors suggest that agrammatic individuals are more impaired in production of regular forms because of their reliance on rule-based procedural memory mechanisms (Ullman et al., 2005), other authors suggest irregular forms are more impaired (de Diego, Costa, Sebastián-Galles, Juncadella, & Caramazza, 2004), and a meta-analysis found no clear dissociation between regular and irregular forms in persons with Broca’s aphasia (Faroqi-Shah, 2007). In summary, the findings surrounding the influence of morphophonological properties of verbs on inflection deficits are variable, and it is unclear if they play a role in sentence production deficits in agrammatism. More information is needed to clarify the nature of verb impairment at the lexical level. Many of these studies also evaluated verb production in a sentence context, intertwining elements of syntactic processing. Finally, none of these accounts consider the function served by verb inflections, and the meaning they convey.

Alternatively, some theories deliberately address verb inflections within their syntactic context. Although agrammatism has been traditionally viewed as a syntactic deficit, several studies found that persons with agrammatism are successful in some syntactic computations, such as determining subject verb agreement (Friedmann & Grodzinsky, 1997), mood marking (Wenzlaff & Clahsen, 2004), and auxiliary verb well-formedness (*will pushed) (Linebarger, Schwartz, & Saffran, 1983). In contrast, other syntactic computations are severely impaired, particularly tense (Faroqi-Shah & Friedman, 2015) and aspect marking (Bastiaanse, 2008; Duman & Bastiaanse, 2009; Fyndanis, Varlokosta, & Tsapkini, 2012). Various authors have explained this observation using the same general idea: when verb inflections fulfil purely syntactic well-formedness functions, as in subject verb agreement, they seem to be spared in production. In contrast, when a verb inflection carries a specific meaning, its production is impaired. Essentially, there is a distinction between “morphosyntactic” and “morphosemantic” processing. “Morphosyntax” refers to the well-formedness constraints of a sentence, while “morphosemantic” refers to the meaning conveyed by a sentence’s morphology.

This general explanation has emerged from multiple different theoretical angles, including syntactic tree structure (the Tree Pruning Hypothesis, Friedmann, & Grodzinsky, 1997) and underspecification of grammatical features necessary for tense marking (Tense Underspecification Hypothesis (TUH), Wenzlaff and Clahsen, 2004). This explanation has also been proposed in the Interpretable Features Hypothesis (Nanousi, Masterson, Druks, & Atkinson, 2006) and the Diacritical Encoding and Retrieval Hypothesis (Faroqi-Shah & Thompson, 2007). Another hypothesis proposes a hierarch of difficulty within tense marking with past tense being more impaired than other tenses (Bastiaanse et al., 2011; Bos, Dragoy, Avrutin, Iskra, & Bastiaanse, 2014), but this was not supported in a meta-analysis (Faroqi-Shah & Friedman, 2015).

Empirical evidence largely supports that semantic implications of verb morphology are a barrier in agrammatic production and comprehension (Faroqi-Shah & Dickey, 2009; Fyndanis et al., 2012). For example, persons with agrammatism are less sensitive to errors in sentences such as “*Last year, my sister lives in Boston”* (Faroqi-Shah & Dickey, 2009). The idea that semantic and interpretable features are impaired in agrammatism has been used to develop treatments for agrammatism (Faroqi-Shah, 2008), which will be discussed in the next section.

The Diacritic Encoding and Retrieval (DER) hypothesis (Faroqi-Shah & Thompson, 2007) suggests that persons with agrammatic aphasia have difficulty with the morphosemantic process that involves matching a temporal adverb marker (such as “yesterday”) to its corresponding verb inflection. Behavioral evidence supports this theory. Faroqi-Shah & Dickey (2009) found that persons with agrammatism are relatively spared in their ability to make grammaticality judgments of morphosyntactic errors, as in the sentence “*The nurse calling a doctor,”* and are most significantly impaired when asked to judge the grammaticality of a morphosemantic violation, as in the example “*Last year, my sister lives in Boston.”* However, this study did not find exclusive support for a morphosemantic deficit; though the impairment was less pronounced, agrammatic individuals were still impaired compared to controls when making morphosyntactic sentence judgments. There was also a single participant in the study who was more impaired in morphosyntactic judgements.

Syntactic-level explanations offer logical theoretical accounts of why persons with agrammatism display syntactic deficits in verb morphology, but they do not directly address the possibility of difficulties with affixation or phonological/lexical impairments in verb inflection production. It is also possible that agrammatism is a reflection of some combination of both levels of impairment; lexical and syntactic deficits need not be mutually exclusive. Thus, further research is necessary to determine whether these deficits occur at the word or sentence level. Furthermore, these explanations do not specify the neural underpinnings of these deficits. We know that agrammatism is linked to lesions in left frontal regions, a region that is also associated with verb impairments (Tyler et al., 2002; Ullman et al., 2005). However, it is not clear what specific neural computations differ between a person with agrammatic aphasia, versus a neurotypical speaker? Do those computations differ during word-level processing tasks, sentence-level processing tasks, or both?

A clear picture has not been formed surrounding the characteristics of verb deficits in agrammatism, and even more specifically, verb tense deficits. Impairment in the ability to properly inflect verbs severely impacts a person’s ability to use language effectively. It is important that we gain a fuller understanding of *why* agrammatic individuals have difficulty in the production of verb inflections, so that we might understand the best routes for remediation of these challenges.

### An intervention for verb deficits and mechanisms of neural plasticity

Given the evidence indicating that a morphosemantic deficit may be at the heart of verb impairments in agrammatism, Faroqi-Shah (2008) designed a “morphosemantic” therapy aimed at targeting these processes. She compared the effects of two interventions for verb impairments, a “morphophonological” therapy targeting oral production in the absence of a sentence context and a “morphosemantic” therapy aimed at improving an agrammatic person’s ability to encode the proper verb inflection given a particular syntactic context. While both therapies resulted in improved aphasia quotients on the Western Aphasia Battery (WAB) in some participants, and a greater variety of inflected verb usage, the morphosemantic treatment resulted in generalization of progress to untrained verbs and improved verb tense accuracy in discourse. The mechanism of this improvement is unclear. Participants may have been improving in their ability to correctly select the appropriate verb inflection for a given context; they also may have been simply improving in their ability to generate correct verb inflections at the word level. Progress was also variable among the participants in this study, who were all characterized as having mild to moderate aphasia (Faroqi-Shah, 2008). While all of the participants showed improved performance at follow-up compared to baseline, maintenance of gains following therapy were varied. Two participants were trained using regular verbs and one was trained using irregular verbs. All three participants showed generalization to untrained regular verbs, however, only two showed generalization to irregular verbs. Faroqi-Shah (2013) then conducted the same morphosemantic treatment with a larger group of participants, and again, participants were trained with either regular or irregular verbs. All participants showed generalization to untrained regular verbs, but no participants showed significant improvement on untrained irregular verbs. Participants also showed mixed improvement in other outcome measures. For example, five participants showed significant improvements on Western Aphasia Battery (WAB) scores, but one did not.

The mixed results surrounding generalization and maintenance are puzzling, and imply that different participants might be using different internal strategies to learn how to use verb inflections. It is possible that those participants who generalize across verb categories improved in their ability to correctly match verb forms to a dictated temporal context, while the participant who did not show the same generalization had learned a simple linguistic rule, such as “add -ed.”

One of the reasons behind the unclear mechanisms for improvement of verb deficits is the considerable lack of neuroimaging data investigating potential neuroplastic effects of therapy. Two prior studies used fMRI to examine the neural activation associated with an intervention targeting syntactic complexity (but not verb tense) for agrammatism, and found increased perfusion and activation in bilateral temporoparietal regions (Thompson et al., 2010; 2013). These were the same regions activated by healthy controls in their experiment, and for several of their participants, constituted perilesional areas. Beyond these two studies, little is known about the neural mechanisms of recovery from agrammatism.

There is considerable research investigating therapy-induced neural plasticity for other symptoms of aphasia, which sheds some light on the mechanisms of language recovery (Hartwigsen & Saur, 2017). Stronger recovery following intervention has been associated with reactivation in left perilesional areas (Breier, Randle, Maher, & Papanicolaou, 2010; Vitali et al., 2007; reviewed in Hartwigsen & Saur, 2017) and arcuate fasciculus (which interconnects temporoparietal and frontal regions) (Breier, Juranek, and Papanicolaou, 2011; Richter, Miltner, & Straube, 2008). Though left hemisphere perilesional activity has been associated with beneficial language processing, the role of the right hemisphere is unclear. The right hemisphere may be either contributing to or interfering with language processing, with some studies showing decreased right hemisphere activation (Richter et al., 2008) and others showing increased activity (Breier et al., 2011; Schlaug, Marchina, & Norton, 2009) associated with recovery. Beyond these spatial localization studies, a deeper understanding of neural plasticity can be gleaned from the handful of studies that have investigated the time course of language processing using ERP or MEG. Neurophysiological responses measured using ERP and MEG have revealed stronger and more distinct components following either word retrieval therapy or intensive language action therapy (Breier et al., 2011; Breier, Maher, Novak, & Papanicolaou, 2006; Cornelissen et al., 2003). When listening to words and pseudowords in a mismatch negativity (MMN) paradigm, the early MMN response (∼50ms) became increasingly left lateralized after intensive aphasia therapy (MacGregor, Difrancesco, Pulvermüller, Shtyrov, & Mohr, 2015). Additionally, during a visual lexical decision task, increasingly strong early ERP components (250-300ms) are found bilaterally in response to real words, but not pseudowords, after intensive therapy (Pulvermüller, Hauk, Zohsel, Neininger, & Mohr, 2005). Word retrieval therapy has also resulted in long latency (300-600ms) activation in the left inferior parietal lobe after therapy when naming pictures (Cornelissen et al., 2003). The time course information from ERP and MEG considerably enhances our understanding of the mechanisms by which perilesional and right homologous regions contribute to aphasia recovery. While MEG has not yet been applied to understanding verb deficits in agrammatism, it could be valuable in revealing the neural mechanisms underlying morphosemantic therapy (Faroqi-Shah, 2008).

### Verb Processing

Before one can begin to characterize the nature of verb deficits and associated neuroplastic changes in agrammatic aphasia, one must understand how verbs are represented in neurologically healthy speakers. It is evident, through a variety of neuroimaging evidence, that verbs and nouns are processed distinctly in the brain by typical speakers (Perani et al., 1999; Shapiro et al., 2005; Tyler, Bright, Fletcher & Stamatakis, 2004); additionally, neuropsychological reports of dissociations in verb and noun processing post brain damage further indicate that they are likely processed separately (Caramazza & Hillis, 1991; Shapiro & Caramazza, 2003). Overall, verbs appear to be represented generally in left frontal and middle temporal regions of the brain; in contrast, nouns tend to be localized in left inferior temporal regions (Perani et al., 1999; Shapiro et al., 2005; Tyler et al., 2004). The left prefrontal cortex has been repeatedly implicated in verb processing specifically (Kielar, Milman, Bonakdarpour, & Thompson, 2011; Shapiro, Moo, & Caramazza, 2006; Shapiro, Pascual-Leone, Mottaghy, Gangitano & Caramazza, 2001). The left frontal regions have been linked to a variety of functions, including verb inflection processing (Kielar et al., 2011; Tyler et al., 2004), selection of competing responses among different verbs (Thompson-Schill, 1998) and complex syntactic processing (Caplan, Alpert, Waters, & Olivieri, 2000; Embick, Marantz, Miyashita, O’Neil, & Sakai, 2000). Temporal lobe activity for verb processing has been associated with both regular and irregular past verbs (Tyler, Stamatakis, Post, Randall, & Marslen-Wilson, 2005).

According to the Full Decomposition Model, prior to any lexical analysis of a word, the brain breaks down a word by its individual morphemes (Taft & Forster, 1975). There is some debate about whether both regular and irregular verb forms are processed through morphological decomposition, or irregular verbs are processed by a separate memory-based system (Marslen-Wilson & Tyler 1997; Sach, Seitz, & Indefrey, 2004; Ullman, 1999; 2004). Regardless, inflected verbs constitute morphologically complex words that are processed fundamentally differently from their uninflected roots. MEG analysis of a lexical decision task has confirmed three distinct temporal stages in complex word processing: recognizing component morphemes, lexical access of individual morphemes, and recombination (Fruchter & Marantz, 2015). The process begins in the left lateral occipitotemporal region at 170ms with visual word recognition (Solomyak & Marantz, 2009). Word processing then follows a posterior to anterior progression, moving to left temporal regions, where lexical access begins (Pylkkänen & Marantz, 2003), and inferior prefrontal regions, with the left inferior prefrontal cortex implicated in processing regular past tense verb inflections (Dhond, Marinkovic, Dale, Witzel, & Halgren, 2003; Fruchter & Marantz, 2015). Some studies have investigated syntactic processing with respect to verb morphology by comparing neurophysiological activity between sentences with and without morphosyntactic violations (Allen, Badecker, & Osterhout, 2003; Kwon et al., 2005; Newman et al., 2007). Morphosyntactic violations result in increased activation in inferior frontal regions and auditory cortices at 300-500ms, and in the middle temporal region at 600ms (Allen, Badecker, & Osterhout, 2003; Kwon et al., 2005; Newman et al., 2007). Regular past tense violations specifically activated left frontal regions at 300-500ms (Kwon et al., 2005; Newman et al., 2007). These findings are summarized in Table 1 and the relevant brain regions are illustrated in Figure 1.

**Table 1:** Summary of time course of verb inflection processing in neurologically healthy speakers. The numbers in parentheses refer to numbers in Figure 1.

| Activity | Tasks | Brain Region | Time | Authors |
| --- | --- | --- | --- | --- |
| Visual orthographic processing (prelexical) | Visual word recognition | Occipitotemporal regions (1) | ~170ms | Solomyak & Marantz, 2009 |
| Semantic, phonemic, and iconic processing of a verb stem | Covert inflection production | Left temporal regions, including Wernicke's area and anterotemporal regions (*see 2, 3 in figure 1*) | ~240ms | Dhond et al., 2003 |
| Lexical access (general) | Visual word recognition | Superior temporal network | ~340-350ms | Solomyak & Marantz, 2009; Pykkänen & Marantz, 2003 |
| Regular and irregular verbs first differentiated | Covert inflection production | Left ventral occipitotemporal lobe (1) |  | Dhond et al., 2003 |
| Detection of morphosyntactic and semantic violations of verb phrases | Sentence reading<br>Sentence grammaticality judgment | Inferior frontal regions (4) and auditory cortices (detection of semantic errors in auditory stimuli) | 400ms (N400) | Kwon et al., 2005 |
| Inflection of regular verbs | Covert inflection production | Left inferior prefrontal cortex (4) | ~470ms | Dhond et al., 2003 |
| Inflection of irregular verbs | Covert inflection production | Right dorsolateral prefrontal cortex (5) | ~570ms | Dhond et al., 2003 |
| Syntactic violations/ grammaticality errors in regular verbs detected | Sentence reading | Middle temporal gyrus (6) | 600ms (N600) | Allen, Badecker, & Osterhout, 2003; |
|  | Sentence grammaticality judgment |  |  | Kwon et al., 2005 |

**Figure 1:**
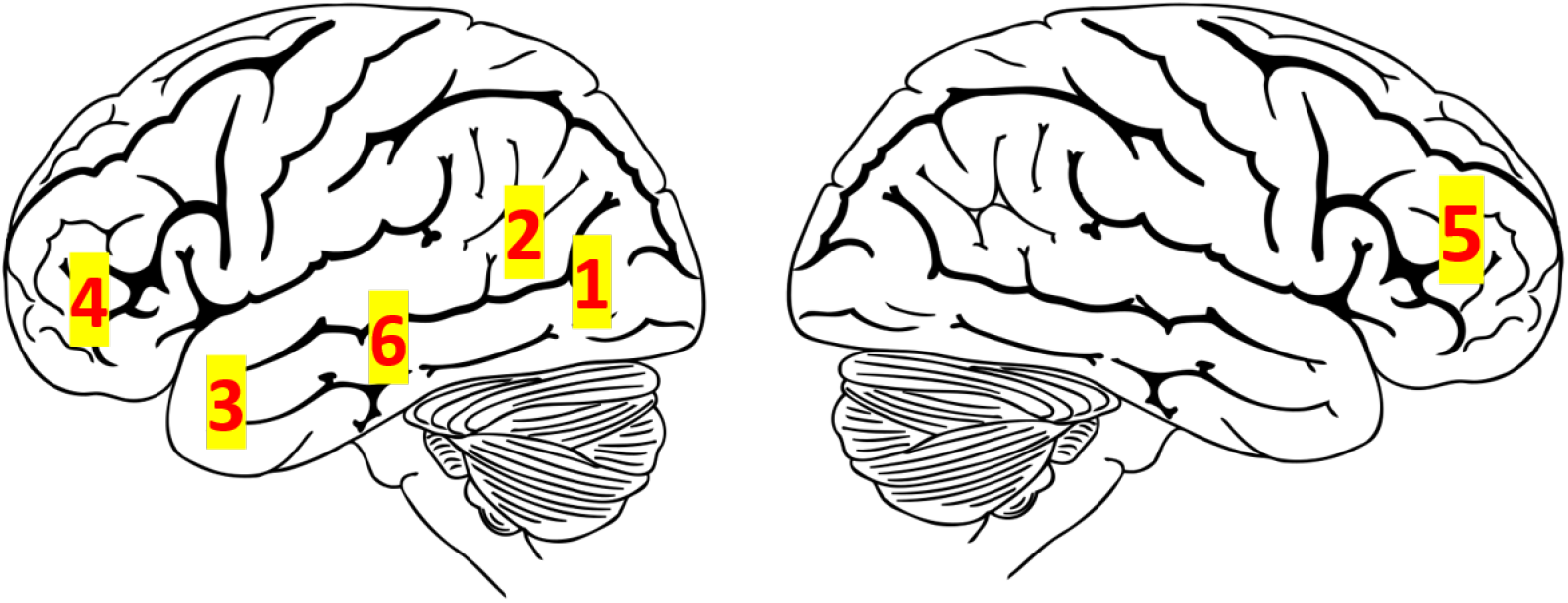
Time course of verb inflection processing in neurologically healthy speakers.

Notably, some of the regions most crucial to verb inflection processing in healthy speakers, such as left inferior frontal regions, are also typically lesioned in persons with aphasia, especially persons with sentence production deficits (Faroqi-Shah et al., 2014). There is a need to further understand these deficits, and to reveal the linguistic level at which they are occurring.

### Summary

This study involved a retrospective data analysis of persons with agrammatic aphasia, who participated in an intervention study and were scanned prior to and following a morphosemantic intervention using MEG. A lexical decision task between inflected and uninflected words (see Table 3), and a sentence judgement task, containing sentences with morphosemantic violations, were used to explore verb processing at the word and sentence level, respectively. Despite the widely held belief that verb deficits, and agrammatism, arise from a deficit in syntactic processing, past behavioral evidence indicates that, additionally, a morphosemantic sentence-level deficit may be at play (Faroqi-Shah & Dickey, 2009; Faroqi-Shah & Thompson, 2007; Fyndanis et al., 2012). Several other studies have explored verb inflection deficits in terms of morphological complexity (de Diego et al., 2004; Ullman et al., 2005), and morphological decomposition (Tyler et al., 2002; Ullman et al., 1997). However, the association between word level and sentence level (morphosemantic) deficits has not been fully explored in the same group of participants.

**Table 2:**
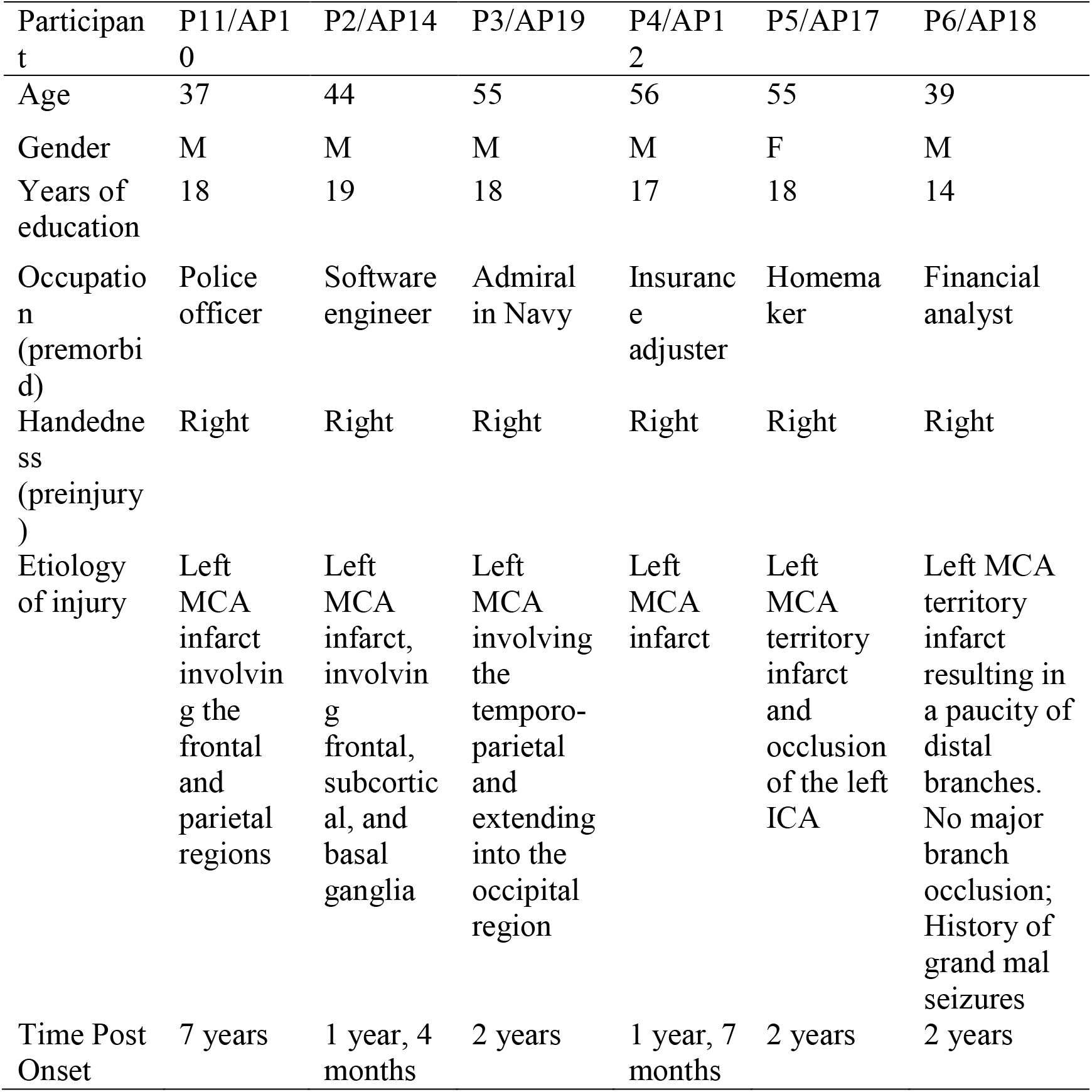
Details of persons with aphasia

**Table 3:**
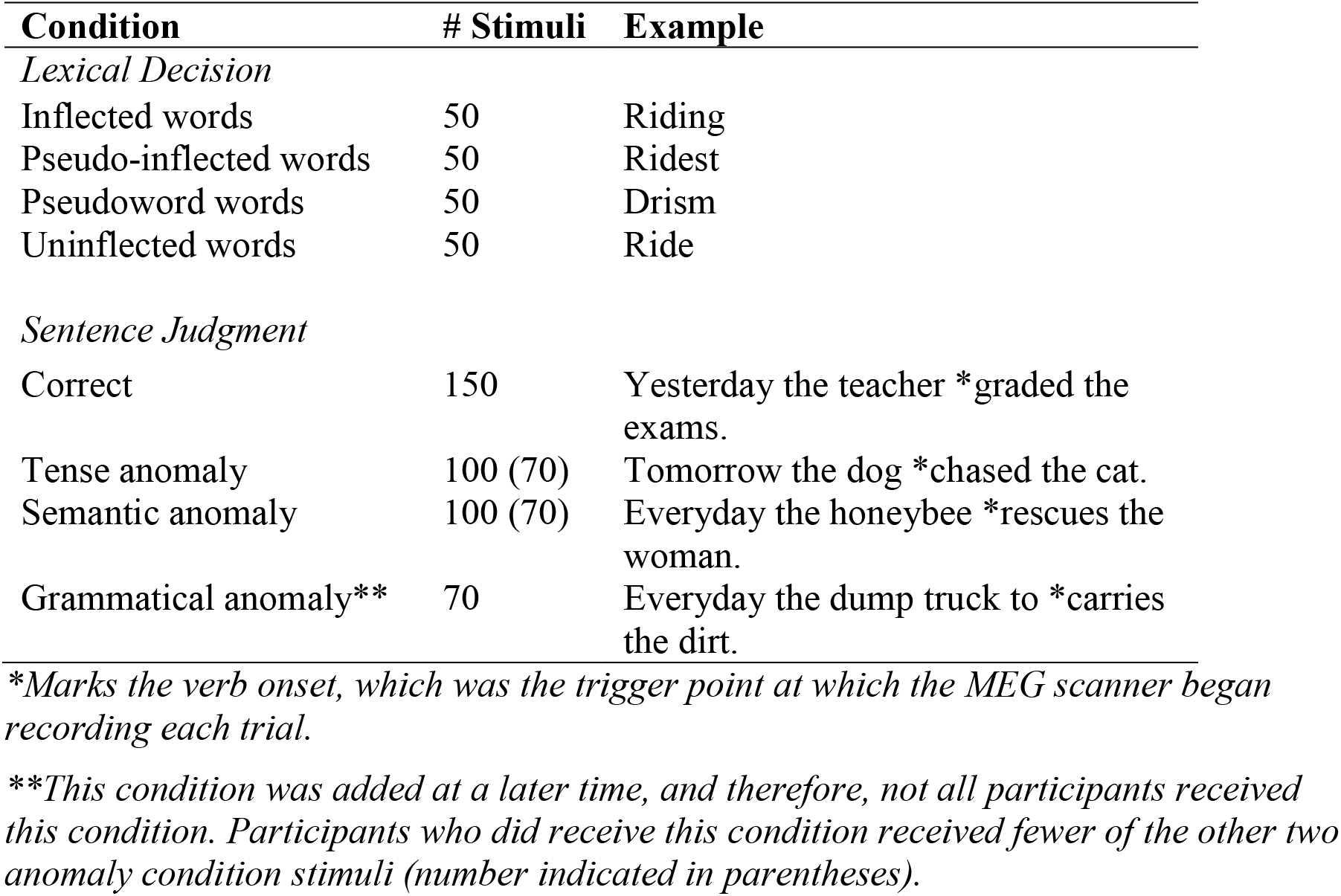
Stimuli for the behavioral judgment tasks

| Condition | # Stimuli | Example |
| --- | --- | --- |
| *Lexical Decision* |  |  |
| Inflected words | 50 | Riding |
| Pseudo-inflected words | 50 | Ridest |
| Pseudoword words | 50 | Drism |
| Uninflected words | 50 | Ride |
| *Sentence Judgment* |  |  |
| Correct | 150 | Yesterday the teacher *graded the exams. |
| Tense anomaly | 100 (70) | Tomorrow the dog *chased the cat. |
| Semantic anomaly | 100 (70) | Everyday the honeybee *rescues the woman. |
| Grammatical anomaly** | 70 | Everyday the dump truck to *carries the dirt. |

*\*Marks the verb onset, which was the trigger point at which the MEG scanner began recording each trial.*
*\*\*This condition was added at a later time, and therefore, not all participants received this condition. Participants who did receive this condition received fewer of the other two anomaly condition stimuli (number indicated in parentheses).*

First, we sought to characterize the nature of verb inflection deficits in agrammatism by comparing patterns of neural activity seen in PWA and controls during the two tasks. We hypothesized that, during the lexical decision task, PWA would show delayed left fronto-temporal responses to stimuli, as past research has demonstrated that M350 peaks in response to single words in PWA tend to be delayed until as late as 600-700ms (Zipse, Kearns, Nicholas, & Marantz, 2011). However, given the behavioral evidence supporting a morphosemantic impairment in agrammatism (Faroqi-Shah & Dickey, 2009), we expected to see the sharpest differentiation between PWA and healthy controls during the sentence judgment task, with PWA displaying relatively decreased and delayed left frontal activation in response to morphosemantic violations.

Second, we sought to determine the linguistic level and neural mechanisms by which verb inflection deficits improve with intervention, and whether or not morphosemantic verb therapy induces neuroplastic changes in the brain. We compared pre- and post-intervention neural activity during both tasks. We hypothesized that the most significant changes in neural activation would be seen post-intervention during the sentence level task, as the morphosemantic intervention specifically targeted verb morphology processing within a sentence context (Faroqi-Shah, 2008; 2013). Given the wealth of recent evidence supporting recruitment of premorbid language areas in aphasia recovery (Thompson et al., 2013; Vitali et al., 2007), we predicted that morphosemantic verb therapy would induce perilesional activation in left frontal regions (Dhond et al., 2003; Kielar et al., 2011; Tyler et al., 2005). Past fMRI data has revealed that aphasia therapy results in faster TTPs in language areas of the brain (Peck et al., 2004; Thompson et al., 2010), therefore, we also predicted that the time course of neural responses to morphosemantic violations in left frontal regions will be faster than in the pre-therapy condition.

## Methods

### Participants

Six persons with agrammatic aphasia (PWA) with a documented verb difficulty were assessed (5 M, 1 F; Mean age of 47.6 years). Additionally, nine age-matched neurologically healthy (AMN) controls (2 M, 7 F; mean age of 69.8 years) and fifteen young neurologically (YN) healthy adults (7 M, 8 F; mean age of 21.6 years) were tested. The AMN and YN groups were pooled together for the purpose of data analysis. All participants were right handed, monolingual, and English-speaking with at least a high school education (mean years of education=17.3) and no significant medical/neurological health concerns or history of psychiatric illness. Participant demographics for PWA are detailed in Table 2, and additional language data from PWA is given in the Appendix.

### Procedures

#### Language testing for persons with aphasia

All participants provided written consent to participate in accordance with procedures set by the Institutional Review Board at the University of Maryland, College Park. The intervention-related behavioral data have been reported in Faroqi-Shah (2008, 2013).

Prior to participating in the present study, all PWA underwent language testing to confirm the aphasia diagnosis and determine a baseline performance level for comparison post-therapy (Faroqi-Shah, 2008; 2013). All PWA were administered the WAB-R (Kertesz, 2006). Narrative speech samples were elicited using pictures and story retelling to generate an overall language profile. Pre-therapy language testing revealed that all PWA had a mild-to-moderate nonfluent (agrammatic) aphasia. Some participants also had difficulty with word repetition. Importantly, PWA had relatively spared auditory comprehension and reading/writing abilities.

Additional other tests were administered to determine proficiency in producing verb inflections in a sentence context, including the verb inflection test (Faroqi-Shah & Thompson, 2004), which evaluates a person’s ability to use verb inflections correctly in a sentence context. Narrative language samples were also specifically analyzed to determine a profile of agrammatism, and revealed diminished sentence structure, limited use of verbs, and mistakes in verb inflections.

Finally, all participants passed a vision screen requiring at least a 20/40 on a Snellen chart (with glasses as necessary) and hearing screen (tested at 500, 1000, and 2000 Hz at 40 dBHL). PWA also passed reading subtest D (single word reading) of the Western Aphasia Battery-Revised (WAB-R) to ensure a high enough reading proficiency to complete the lexical decision task (Kertesz, 1982). (Note—Participant P3 did not complete subtest D of the WAB, but did pass subtest A, sentence reading).

#### Language intervention for persons with aphasia

Persons with aphasia completed a morphosemantic therapy, which emphasized improving a person’s ability to associate a temporal context with a verb inflection within a sentence. The morphosemantic therapy for verb inflection deficits was administered using either regular or irregular verbs as stimuli. Twenty verbs were trained directly during therapy, and 20 were used to probe for generalization. Irregular verbs consisted of either a verb with a vowel change, or a vowel change and a consonant addition. Participants were randomly or pseudo-randomly assigned to either regular or irregular verb conditions in both Faroqi-Shah (2008) and Faroqi-Shah (2013). Participants completed three phases: baseline testing, treatment, and follow-up testing. Treatment was administered 4-5 days/week in 1-2 hour long sessions. Treatment continued until a participant reached 90% accuracy on treatment probes 4 sessions in a row, after 20 sessions, or after demonstrating a failure to improve by more than 10% after 12 sessions.

The therapy included 5 steps: confrontation naming: naming an action depicted in an image; anomaly judgement: identifying mismatches in adverbs and verb tenses in sentences; auditory comprehension: matching a presented sentence to a picture, from a choice of three representing three different verb tenses; sentence completion: writing down a correctly inflected verb for a sentence corresponding to a picture; and sentence construction: arranging written word cards in the correct order to form a sentence. Each step was completed with each verb. Training emphasized verb inflection usage within a sentence context, and oral production was discouraged during this therapy, in order to isolate improvement due to morphosemantic, rather than morphophonological improvements. On average, participants took 15 treatment sessions to complete the therapy, with an average of over 90% accuracy for trained verbs upon completion.

### Neuroimaging

All of the participants received whole-head MEG scans during completion of two behavioral judgment tasks, a lexical decision task and a sentence judgment task (described below). MEG scans were conducted at the University of Maryland, College Park using a 160-channel MEG scanner (Kanazawa Institute of Technology (KIT), Japan). Head coordinates were calculated using the Polhemus 3SPACE FASTRAK system. Neuromagnetic fields were recorded using a 160-channel axial gradiometer whole-head system, sampled at a rate of 500 Hz and bandpass filtered at 1-200 Hz. During each experimental block, event-related neuromagnetic responses were measured, averaged, filtered at 40 Hz lowpass, and adjusted with a 100ms pre-stimulus baseline. Scanning for the lexical decision and sentence judgment tasks happened on the same day.

### Behavioral Judgment Tasks for MEG

Experimental stimuli were presented using version 8.0 Presentation software (Neurobehavioral Systems). Practice trials were presented before experimental trials. Stimuli for the lexical decision task were presented visually, while stimuli for the sentence judgment task were presented auditorily. Visual stimuli were presented onto a monitor located in the MEG scanner booth in 128-point Times New Roman lower-case font. Auditory stimuli were presented with external speakers at a comfortable listening level. Participants received verbal instructions from an experimenter instructing them on which keys to press.

#### Lexical decision task

This task, which was designed to examine verb inflection processing at the word level, had four conditions: inflected words, pseudo-inflected words, pseudoword words, uninflected words (Table 3). Stimuli for each condition were developed from regular English verbs with high usage frequency. Verb inflections were regular English verbs that ended in either -d, -s, or -ing. “Pseudo-inflections” were generated with the suffixes - al, -ly, or -en affixed onto verb stems. Pseudowords were created by switching one letter in the word (as in survive-surpive). The four experimental conditions created an orthogonal comparison between real words (verb stems and verb inflections) and nonwords (pseudonflections and pseudowords) as well as morphologically simple (verb stems, pseudowords) and complex (verb inflections and pseudo-inflections) words.

Stimuli (words) were presented to the participants in a pseudorandom order, for 500ms, with 3000ms to respond before the next stimulus appeared. Participants were told to decide if the stimulus presented was a real word or not, and press a button with their left hand (to account for right-sided weakness due to CVA) to indicate their choice. They did not receive feedback about the correctness of their response. Task stimuli were presented in four experimental blocks of two minutes each, with 30 seconds in between.

#### Sentence judgement task

This task, which was designed to examine verb inflection processing in a sentence context, had four sentence conditions: correct, tense anomalies, semantic anomalies, and grammatical anomalies (Table 3). In the tense anomaly condition, sentences were created using real words that were either correctly or incorrectly matched to temporal markers (i.e., “yesterday”) in a sentence. Stimuli were varied so that the adverb sometimes occurred before the verb and sometimes after the verb. The location of the tense marker (in an auxiliary verb or as part of a main verb) and the type of tense (past, present, or future) were also varied across stimuli. The crucial contrast was between correct sentences and tense anomaly sentences, which required a nuanced judgment about the suitability of the verb inflection given a temporal context. This construct was an important focus of the morphosemantic intervention (Faroqi-Shah, 2008; 2013). A semantic anomaly condition was also included, because past research has demonstrated that persons with agrammatism have relatively spared semantic processing abilities (Faroqi-Shah & Dickey, 2009). In the semantic anomaly condition, a grammatical sentence was constructed using words that were unfitting in meaning. Finally, a fourth condition, a grammatical anomaly, was included for some participants, in order to differentiate morphosemantic processing from general syntactic processing.

Stimuli were audio recorded by a male speaker of English. The sound waves of each sentence were used to note the time of verb onset using Praat software (Boersma & Weenik, 2004). These verb onset times were used to set the MEG trigger. During the experimental procedures, sentences were presented auditorily in a pseudorandom sequence. Each trial began with an auditory beep. The sentence was played 300ms later, and the next trial began 7500ms after the onset of the prior trial. A more detailed description of the sentence judgment task, and the detailed results of participant’s performance on this task (outside the scanner) pre-intervention, are reported in Faroqi-Shah and Dickey (2009). Participants were asked to make a judgment about whether or not the sentence was “correct” or “incorrect,” by pressing one key for “correct” and another for “incorrect.” Participants again responded using their left hand to account for potential right side hemiparesis. No feedback surrounding the accuracy of responses was given.

### Behavioral Data Analysis

Accuracy and reaction time information was collected for controls and PWA during both experimental tasks and recorded in individual log files. This data was collected and analyzed in SPSS software version 24 (SPSS, 2016). Accuracies were converted to rationalized arc sine units (Studebaker, 1985).

For the lexical decision task, accuracies in RAU were compared between groups and across conditions using a univariate General Linear Model. Accuracy data for the sentence judgment task was not analyzed due to experimental error. Reaction times for both tasks were compared using generalized linear mixed effects model (GLMM, Hedeker, 2005) on raw (untransformed) response times (Lo & Andrews, 2015) with group (controls and PWA), task conditions, and the interaction term as fixed factors, and items and participants as the random factors. The intercept was included in the model for fixed and random factors.

### MEG Data Analysis

#### Preprocessing

Data analysis of MEG scans was conducted using MNE Suite Software (Gramfort et al., 2014) using the Eelbrain pipeline (Brodbeck, 2019). Original MEG raw files were bandpass filtered at 0-40 Hz in order to reduce external noise and remove artifacts that resulted from ocular or muscular movements. Unresponsive and noisy channels (identified by MNE Suite or via visual inspection of raw data) for individual participants were also excluded from data analysis. Due to excess noise in MEG scans from the Age Matched Normal (AMN) group, independent component analysis (ICA) was computed for all participants, to further reduce the impact of external artifacts on the raw data. ICA is a statistical procedure that identifies and extracts independent source signals, allowing for subsequent removal of sources originating from external artifacts, while retaining sources originating from neural signals (Gramfort et al., 2014; Vigário, Sarela, Jousmiki, Hamalainen, & Oja, 2000). Once ICA was calculated for each participant, components were visually inspected and components resulting from eye blinks, heartbeats, and other non-neural signals were removed.

Within each task, trials were sorted by experimental condition, and epoched from -100 to 800 and 1000ms time windows for the lexical decision and sentence judgment tasks, respectively. For the lexical decision task, epochs began at the word onset; for sentence judgment tasks, epochs began at the time of the verb onset (verb onset is marked with a * in Table 3). Time windows of 800 and 1000ms were selected to capture neural activation in PWA, who may have delayed response times to stimuli compared to neurologically healthy speakers, who produce typical responses at about 400ms for single words (Pylkkänen & Marantz, 2003) and up to 600ms for sentences (Allen, Badecker, & Osterhout, 2003) (see also Table 1). Individual trials were visually inspected, and trials with excessive noise were removed from further averaging. At least 80% of the collected trials were retained for each participant after removal of bad trials. Finally, responses from all trials of a given condition were averaged (combined) together for source estimation.

#### Source Estimation

MEG scans for all participants were overlaid with a standard structural MRIs in order to compute more specific source localization. An MRI template, *FSaverage,* supplied by FreeSurfer, was used (Fischl, Sereno, Tootell, & Dale, 1999). *FSaverage* was co-registered to the participant’s head shape based on the location of MEG sensor markers. During co-registration, the iterative closest point (ICP) was established by converting MEG digitizer points to MRI coordinates, and finding the alignment where digitizer points were most closely matched to head surface coordinates. Source estimates were calculated using the minimum norm estimate approach (Hämäläinen & Ilmoniemi, 1994). In order to restrict our search for effects to specific brain regions, parcellations of regions of interest were defined based on the division described in Desikan et al. (2006).

Data analysis was conducted using a combined “control” group of age-matched normal (AMN) and young neurologically healthy adults (YN). During the pre-processing stage, it was noted that AMN data was particularly noisy, and after removing artifacts originating from external sources, neural signals from this group could not be detected. Conducting initial data analyses on the two control groups separately revealed that the signal was only detectable when both groups were included.

#### Statistics

To avoid the high probability of type 1 error when comparing many MEG sensors at multiple time points, nonparametric cluster based permutation tests were run instead of parametric statistical tests, as described in Maris & Oostenveld (2007). For each task and within each group, cluster-based 1-way ANOVAs were run, with condition as the independent variable. Subsequent post-hoc related measures t-tests were run to further determine sources of main effects observed (Brodbeck, 2019).

MEG responses for the lexical decision task were statistically analyzed using separate one-way ANOVAs for each group, with the word conditions as the independent variable (four levels—real-inflected, real-uninflected, pseudo-inflected, and pseudoword). Based on the brain response patterns outlined in Table 1, for healthy controls, the MEG responses were analyzed in two separate time bins and brain regions. Responses in the left fusiform gyrus were analyzed from 150ms to 300ms, to look for neural effects of early, orthographic-based decomposition of inflected words, given that early morphological decomposition has been associated with left occipitotemporal regions (Fruchter & Marantz, 2015; Lehtonen, Monahan, & Poeppel, 2011; Morris & Stockall, 2011). The other test was run from 300-500ms, in a left frontotemporal parcellation. This test was conducted to look for the effects of accessing lexical (semantic) meaning, typically evidenced by the well-documented N400 effect (Pylkkänen & Marantz, 2003), and later decomposition of stems and inflections (Dhond et al., 2003; Fruchter & Marantz, 2015). Corresponding post-hoc t-tests were conducted when significant effects were found.

For PWA, the same statistical tests were completed, twice—for pre-intervention and post-intervention MEG scans. Tests were conducted similarly, though time bins for PWA were expanded slightly (150-400ms, 300-800ms) to account for delayed word processing in aphasia (Zipse et al., 2011).

For the sentence judgment task, a one-way ANOVA with violation type as the independent variable (three levels—no violation, tense violation, semantic violation) was conducted. Because not all participants had completed the “grammatical” violation condition, this condition was not analyzed in the present study. This test was conducted in two separate time bins—one from 300-500ms, and one from 500-700ms, to look for the effects of early and late grammatical processing, as described in Kwon et al. (2005), and Table 1. Consistent with Kwon’s findings, searches were restricted to left frontal and temporal regions.

Finally, for both tasks, tests were also run in corresponding right hemisphere regions for PWA, to account for the possibility of right hemisphere compensation of language skills (Kinsbourne, 1998). Again, time bins were expanded in PWA to account for delayed processing. If significant right hemisphere effects were found in PWA for a particular test, it was replicated in healthy controls to differentiate between compensation and pre-existing normal activity.

## Results

### Behavioral Results

#### Lexical decision

##### Accuracy

When comparing PWA to controls, there was a main effect of group (F=20.7, p<.001) and condition (F=6.6, p<.001) and a significant group x condition interaction (F=5.3, p<.01). Planned pairwise comparisons using Mann-Whitney U test showed that PWA were less accurate than healthy controls for uninflected (Mann-Whitney U test for independent samples, U = 6, p<.01), inflected (Mann-Whitney U = 14.5, p<.05) and pseudoinflected (Mann-Whitney U = 3, p<.01) words. For PWA, there was no difference between overall accuracy across conditions before and after treatment (Related Samples sign test, z = 1, p=.3)

##### Reaction time

When comparing controls to PWA pre-intervention, there was a main effect of group (F = 9.5, p <.001) and word type (F =62.5, p < .001). In addition, there was a significant interaction between group and word type (F=17.7 p < .001). That is, persons with aphasia (Mean RT = 1.44 seconds) were slower than controls (Mean RT = 1.04 seconds). The interaction showed that the two groups had a slightly different response pattern across word types. Both groups responded more slowly to morphologically complex words (inflected and pseudo-inflected) compared to simple words (uninflected and pseudowords). While controls showed no RT difference between inflected and pseudo-inflected words, PWA were significantly slower for pseudo-inflected words. Within simple words, controls were significantly slower for pseudowords compared to uninflected words, while PWA showed no difference between uninflected and pseudowords.

Post-intervention, there were still main effects for group (F=18.01, p<0.001), with PWA reacting more slowly than controls, and condition (F=71.50, p<0.001), with morphologically complex words eliciting slower reaction times. Interactions between group and condition also remained post-intervention (F=24.29, p<0.001), with PWA performing more slowly for pseudo-inflected words and showing no difference in RT between pseudowords and uninflected words.

#### Sentence Judgment

Due to experimental error, accuracy data was not reported for the sentence judgment task, but reaction time data was collected and analyzed. Reaction times for the tense violation, semantic violation, and correct sentence conditions were analyzed, excluding those above 5000ms or below 300ms. Reaction times were compared using generalized linear mixed effects model (GLMM, Hedeker, 2005) on raw (untransformed) response times (Lo & Andrews, 2015) using SPSS version 24 (SPSS, 2016) with group (control, aphasia), sentence category (tense violation, semantic violation, and correct sentences) and the interaction term as fixed factors, and items and participants as the random factors. The intercept was included in the model for fixed and random factors.

Pre-treatment, when comparing PWA to controls, there was a main effect of condition (F=18.94, p<0.001), with reaction times for the tense violation condition being longer than the semantic violation or correct sentence conditions, and a main effect for group (F=11.35, p=0.01), with PWA having slower reaction times (PWA mean RT=2.27 seconds, controls mean RT=1.79 seconds). In addition, there was a significant interaction between group and condition, (F=8.35, p<0.001), which revealed that persons with aphasia were fastest in their response to the semantic violation condition, but controls were fastest in their response to correct sentences. When comparing reaction times for PWA pre-intervention to post-intervention, there was a main effect for time (F=11.21, p=0.001) and condition (F=54.70, p<0.001), and an interaction, such that persons with aphasia showed slower reaction times in response to the tense violation and control conditions post-intervention, but no change was seen in response to the semantic violation condition (F=4.34, p<0.013). Comparisons of reaction times of PWA post-intervention to controls revealed the same patterns as pre-intervention comparisons.

### MEG Results

#### Lexical decision

In the left fusiform gyrus region, neurologically healthy participants showed a main effect of condition in the 220-300 ms time range (F_max_=6.4, p=0.03). Planned pairwise post-hoc t-tests revealed that the MEG response differed between the pseudoinflected words and pseudowords (t_max_=0.042, p=0.04) (Dale et al., 2000). Analysis of the 300-500 ms time range in the frontotemporal regions revealed a main effect for condition. The effect was centered in left frontal regions and lasted for 130ms, between 300-430ms (F_max_ =5.32, p=0.003). Planned pairwise post-hoc t-tests also revealed significant differences between specific conditions (Figure 3). One effect occurred between real-inflected and pseudo-inflected conditions. The effect lasted for 110ms, between 390-500ms, and was centered in the left temporal cortex, (t_max_=2.85, p=0.04). Another occurred between real-inflected and pseudoword words, lasting for 120ms, between 300-420ms. This one was centered in the left frontal cortex (t_max_=3.51, p=0.02).

**Figure 2:**
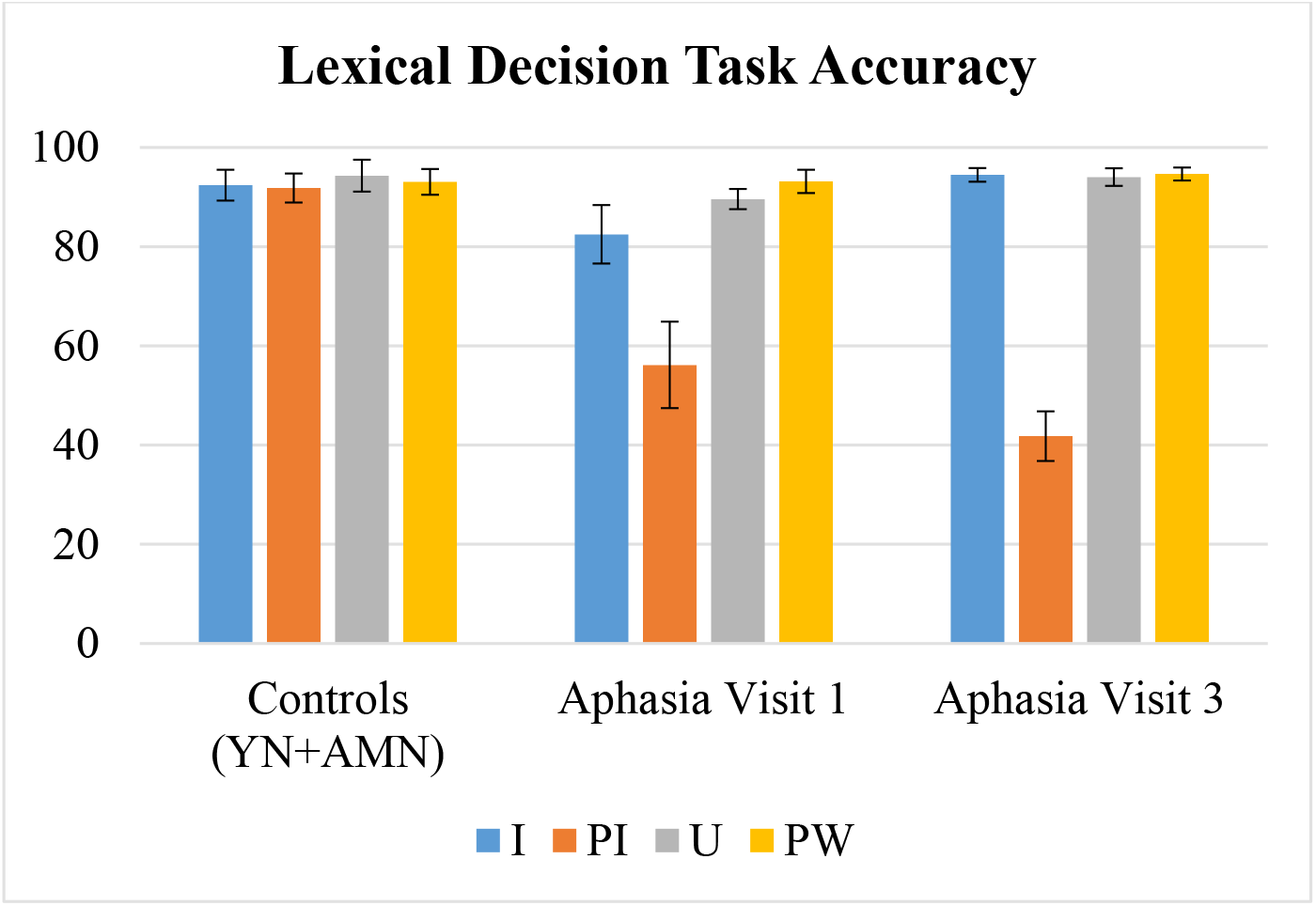
Lexical decision accuracy results. I=inflected, PI=pseudo-inflected, U=uninflected, PW=pseudoword.

**Figure 3:**
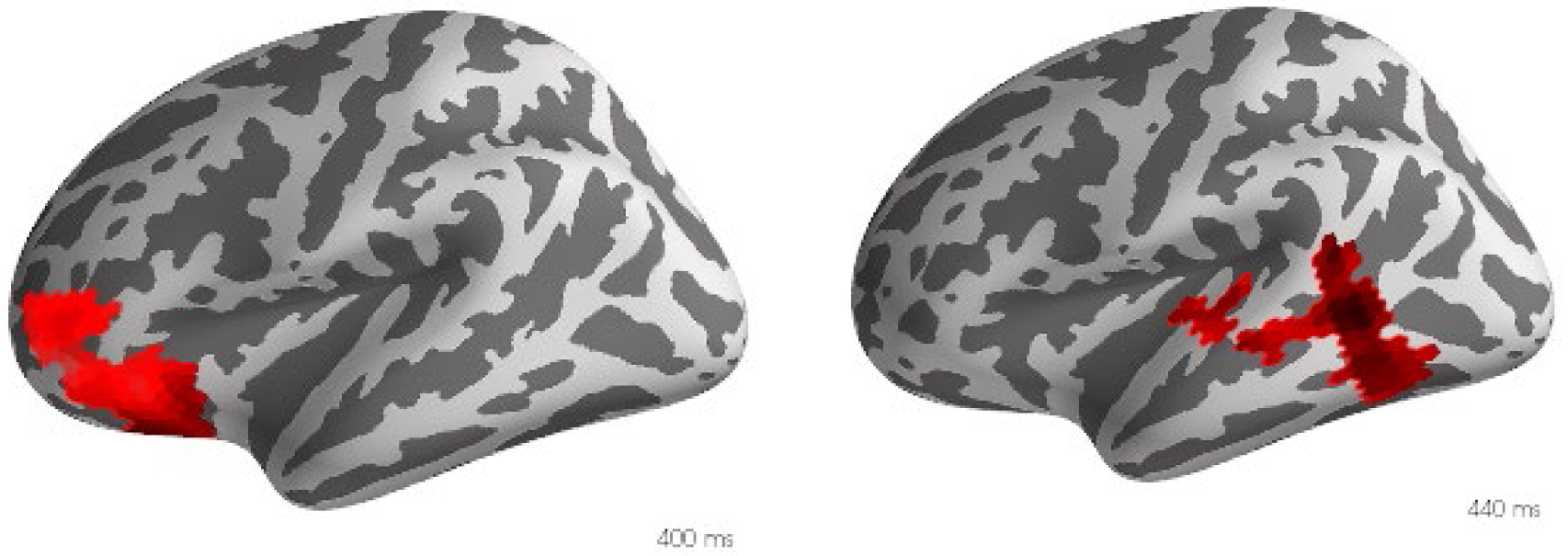
MEG response to lexical decision task in healthy controls. a) Real-inflected words> pseudowords, left frontal, 400ms, p=0.04 b) Real inflected words>pseudo-inflected words, left temporal, 440ms, p=0.02.

Pre-intervention, persons with aphasia did not show sensitivity to any of the different conditions for the lexical decision task in this early time bin (expanded from 150-400ms), nor did they show differences among conditions in the later time bin (expanded from 300-800ms). Post-intervention, again, no significant differences across conditions were noted for the lexical decision task, in either time bin.

##### Right hemisphere results

PWA did not show sensitivity among different word-level conditions in a 1-way ANOVA in the right fusiform gyrus. Across a right hemisphere frontotemporal parcellation, PWA did not show sensitivity among different conditions in the lexical decision task in right frontotemporal regions pre- or post-intervention.

#### Sentence Judgment

For the sentence judgment task, effects were investigated in left frontal and left temporal regions in two time bins—300-500ms, and 500-700ms. Control participants showed significant effects in both frontal and temporal regions in the earlier time range. From 360-470ms, there was a strong main effect in the left frontal cortex (F_max_=10.33, p=0.001), and from 300-500ms, there was a significant main effect in the left temporal cortex (F_max_=16.96, p=0.006). Subsequent post-hoc t-tests revealed that results were significant for the “semantic>tense” contrast in the left frontal regions between 360-490ms (t_max_=4.78, p=0.004), and the “semantic>tense” and “semantic>none” conditions in the left temporal lobe, between the 300-500ms and 410-470ms ranges, respectively (t_max_=5.6, p=0.0004 and t_max_=6.23, p=0.02). For the later time bin (500-700ms), a significant main effect (though not as strong), was found in left temporal regions between 500-550ms (F_max_=9.16, p=0.048). Subsequent post-hoc t-tests revealed this effect to be originating from the “semantic>tense” condition between 500-570ms (t_max_=4.52, p=0.049).

The same statistical tests were computed for PWA’s pre-intervention therapy scans. Despite expanding the time bins to account for delayed processing in PWA, no significant differences were found among conditions in the pre-intervention sentence judgment scans. However, significant differences were noted for the grammaticality judgment task in the late time bin (500-1000ms for PWA, because the time bins were expanded) post-intervention. A one-way ANOVA between the different sentence conditions revealed a significant effect in the left temporal lobe between 700 and 770ms (F_max_=17.62, p=0.03), and subsequent post-hoc t-tests revealed that the effect was originating from the “semantic=tense” violation condition, between 720-750ms (t_max_=5.02, p=0.03).

##### Right hemisphere results

Pre-intervention, PWA did not show sensitivity among sentence judgment task conditions in the right frontal or right temporal lobe in either time bin, with one exception. There was a strong effect for the semantic=tense condition in the right temporal region *pre-intervention* between 310-391ms (t_max_=8.45, p<0.001). When the same test was run in healthy controls (300-500ms time bin), they also showed a significant effect, between 300-480ms (t_max_=5.41, p=0.003). Interestingly, this right temporal response was *not* present post-intervention in PWA.

Post-intervention, PWA only showed one significant response to sentence judgment tasks in the right hemisphere: a late right temporal lobe response post-intervention to the “semantic>tense” condition from 960-1000ms (t_max_=7.41, p=0.03). Follow up testing confirmed that healthy controls also showed this response, from 940-1000ms (t_max_=3.81, p=0.03), but PWA did not show it pre-intervention (t_max_=6.01, p=0.73).

Given these patterns, the answer to the first research question is that PWA are impaired in processing verb morphology at the word level. No significant differences were noted among conditions for the word-level task pre-intervention, and minimal significant responses were assessed for the sentence-level task pre-intervention. Sentence-level deficits may be downstream effects, as word-level processing is necessary for sentence-level tasks. However, PWA did show significant differences in their neural responses to different conditions in the sentence-level task post-intervention. Thus, the answer to the second research question is that improvements in verb morphology production post-intervention occur via changes in sentence-level processing.

## Discussion

The purpose of this study was to a) determine the nature of verb inflection deficits in agrammatic aphasia and b) determine the mechanism by which these deficits are remediated with intervention. Analysis of MEG brain responses collected during a lexical decision (word-level) and grammaticality judgment (sentence-level) task reveals that persons with agrammatic aphasia are likely experiencing deficits at both the word and sentence level, but that training-induced improvement in verb inflection processing occurs at the sentence level.

### Characterizing Verb Morphology Deficits

#### Word-level processing

Our first research question considered the nature of verb inflection impairment in agrammatism, and the level at which this breakdown occurred. We first analyzed the results of the lexical decision task. Our lexical decision task investigated word-level verb inflection processing with four conditions: real-inflected, real-uninflected, pseudo-inflected, and pseudoword. The pseudo-inflected condition consisted of real word stems with illegal inflections (“ridest”), while the pseudoword condition consisted of unreal word stems (“drism”). Thus, sensitivity to the pseudo-inflected condition required morphological decomposition in order to determine that two legal components (“ride” + “est”) were not permitted together, while sensitivity to the pseudoword condition required sensitivity to the lexical (semantic) status of a word.

The lexical decision task was presented as a visual paradigm to ensure that the morphological composition of inflected words was clear—coarticulation occasionally decreases the salience of consonant clusters over audio, and the morphological unnaturalness of pseudo-inflections may not have been evident with auditory presentation. Analysis of the behavioral data for the lexical decision task revealed that PWA were less accurate in their decision-making compared to healthy controls (Figure 2) and displayed slower reaction times. There was also no significant change in accuracy pre- to post-intervention for PWA, though we only looked at overall changes in accuracy, and did not analyze pre- to post-change in individual conditions. So, it is possible that there may have been accuracy improvement in individual conditions. Additionally, the stimuli from the lexical decision task included repetition of the same word stems (e.g., “ride,” “riding,” “ridest,” see Table 3). Because the same stems were presented multiple times, it is possible that semantic priming (Bentin, McCarthy, & Wood, 1985) or repetition effects (Ratcliff, Hockley, & McKoon, 1985) may have influenced decision-making in controls and PWA (Milberg & Blumstein, 1981).

Analysis of neuroimaging data revealed that healthy controls were sensitive to both the morphological structure of those words (real-inflected versus pseudo-inflected) and the lexical status of a word (real-inflected versus pseudoword) (see Figure 3). Controls showed sensitivity to differences among different conditions in the lexical decision task between 220-300ms, in the left fusiform gyrus (Table 4a), an area previously associated with morphological decomposition of inflected words (Lehtonen, Monahan, & Poeppel, 2011). This sensitivity was found to be between the pseudo-inflected and pseudoword condition, indicating a neural response to differences in structure among nonwords. This effect is somewhat consistent with other reports of the N250, an ERP component revealing early morphological parsing (Morris & Stockall, 2011), and other reports that have found occipitotemporal activity reflective of representations of morphological structure at about 220ms (Lehtonen, Monahan, & Poeppel, 2011). However, this finding is inconsistent with accounts of the M170 effect, showing very early sensitivity to morphological structure (Fruchter & Marantz, 2015). It is also puzzling that our effect was only seen in pseudo-words, indicating a rapid parsing for words that one has never seen before, but not for familiar words. Prior literature has indicated that the process of morphological decomposition occurs regardless of lexical features, including word frequency (McCormick, Brysbaert, & Rastle, 2009; Rastle, Davis, & New, 2004), so we would have expected no difference in inflection parsing between real and psuedowords. Our results are possibly supportive of a dual-route processing model, whereby some complex words are recognized via a whole word mechanism that does not rely on decomposition, but pseudowords, which do not have a whole word entry in a mental lexicon, must be decomposed (Vannest, Polk, & Lewis, 2005).

**Table 4a:** Summary of Lexical Decision Results showing pairwise contrasts, 150-300ms, *NS = not significant, I= inflected, U=uninflected, PU=pseudowords, PU=pseudouninflected*

| Controls |  |  |  |  | PWA (Pre-intervention) |  |  |  | PWA (Post intervention) |  |  |  |
| --- | --- | --- | --- | --- | --- | --- | --- | --- | --- | --- | --- | --- |
| *Left fusiform gyrus* |  |  |  |  |  |  |  |  |  |  |  |  |
|  | I | U | P | PI | I | U | P | PI | I | U | P | PI |
| I | - | - | - | - | - | - | - | - | - | - | - | - |
| U | NS | - | - | - | NS | - | - | - | NS | - | - | - |
| P | NS | NS | - | - | NS | NS | - | - | NS | NS | - | - |
| PI | NS | NS | p=0.04 | - | NS | NS | NS | - | NS | NS | NS | - |

Controls also showed sensitivity to the morphological structure of pseudo-inflected words in the left frontal lobe between 390-500ms (Table 4b), indicating a later sensitivity to morphological complexity. This later effect may be reflective of recombination of components (“ride” + “ing” = “riding”) (Taft, 2004), which Fruchter & Marantz (2015) have theorized occurs between 430-500ms. The effect was also seen in the left temporal lobe, and may be indicative of the N400 effect, because successful decomposition of our pseudo-inflected words should have alerted the participant to the nonword status of the stimulus.

**Table 4b:** Summary of Lexical Decision Results showing pairwise contrasts, 300-500ms

| Controls |  |  |  |  | PWA (Pre-intervention) |  |  |  | PWA (Post intervention) |  |  |  |
| --- | --- | --- | --- | --- | --- | --- | --- | --- | --- | --- | --- | --- |
| *Left frontotemporal regions* |  |  |  |  |  |  |  |  |  |  |  |  |
|  | I | U | P | PI | I | U | P | PI | I | U | P | PI |
| I | - | - | - | - | - | - | - | - | - | - | - | - |
| U | NS | - | - | - | NS | - | - | - | NS | - | - | - |
| P | p=0.02 | NS | - | - | NS | NS | - | - | NS | NS | - | - |
| PI | p=0.04 | NS | NS | - | NS | NS | NS | - | NS | NS | NS | - |

Given that controls showed significantly different responses to the real-inflected vs pseudo-inflected and real-inflected versus pseudoword conditions in the 300-500ms time bin, our results are consistent with past findings that a word’s lexical status (real or not real) is retrieved in this time range in left frontotemporal regions (Pylkkänen & Marantz, 2003; Solomyak & Marantz, 2009). Other MEG studies investigating single word processing have also asserted that lexical meaning first comes into play in visual word recognition around 300ms (Solomyak & Marantz, 2009), and several have documented the M350 component—a frequency-mediated neural response reflecting lexical-semantic processing of a word (Embick, Hackl, Schaeffer, Kelepir, & Marantz, 2001; Stockall, Stringfellow, & Marantz 2004). Future investigations of complex, single-word processing in agrammatism might consider the impacts of word frequency on verb inflection processing.

Despite the expected patterns of neural activation seen in healthy controls, persons with aphasia showed no significant responses to morphological complexity or lexical status differences among conditions in the lexical decision task. Word level processing is inherently a precursor to sentence level processing, so this data indicates that part of the verb morphology impairment seen in agrammatism includes a verb inflection deficit occurring at the word level. These results are somewhat surprising, and it is possible that our small sample size for PWA (n=6) resulted in decreased statistical power to detect a response. However, given a neural response to the sentence level task post-intervention was measured, this is not the most likely interpretation. It appears that participants had difficulty with word level processing of morphologically complex words. This is surprising and novel finding, as agrammatism has classically been described as a syntactic deficit (Berndt & Caramazza, 1980; Friedmann, 2001). However, Badecker (1997) describes one case study of a person with aphasia whose deficits in grammar originate from morphological errors at the word level, including inflection deletion and affix substitutions. Because word-level processing and decomposition are inherently involved in sentence processing, a word level deficit would have also contributed to the lack of response seen in our PWA to the sentence-level grammaticality judgment task pre-intervention.

Notably, though persons with aphasia did not show neural sensitivity to differences among conditions in the word-level task, our behavioral results indicate that they were capable of differentiating real versus pseudowords in practice, as they demonstrated accuracies greater than 80% for three out of four lexical decision conditions (Figure 2). Thus, the neural processing behind the lexical decision task must be occurring via a compensatory route that our methods did not expose. Because we based our search for effects during the lexical decision task on what was seen in healthy controls, PWA may be relying on regions not typically devoted to language processing during the lexical decision task.

A word level impairment also points to the possibility that non-syntactic factors are coming into play when people with agrammatism process verbs. Early accounts of agrammatism suggested that Broca’s aphasia is actually the result of a word level phonological deficit, which often contributes to the omission of bound morphemes and function words (Kean, 1977). More recent accounts have branched from the idea of a solely phonological deficit, citing morphophonological parsing as the source of the deficit in persons with aphasia (Tyler, Randall, & Marslen-Wilson, 2002). Our patients did not show sensitivity to differences among inflected and uninflected conditions, pointing to the possibility that morphophonological parsing may indeed have been a source of difficulty. Future research could investigate whether this parsing difficulty is unique to verbs, as morphologically complex verbs have been found to be uniquely represented in the brain (Tyler, 2004), or if this impairment is seen in all inflected structures.

#### Sentence-level processing

We then examined the results of the sentence judgment task. Analysis of the behavioral data revealed that PWA were slower in their responses than healthy controls, and that all participants were slower in their responses to sentences with tense violations. The sentence judgment task was presented auditorily, unlike the lexical decision task, which was visual. As mentioned above, the lexical decision task was presented visually to ensure detection of less salient verb inflections. We acknowledge that decreased auditory salience of verb inflections may have also influenced perception during the sentence judgment task, especially when the only marker of tense in a sentence was a word-final inflection, given that final consonants are typically less perceptually distinctive (Redford & Diehl, 1999).

Kwon et al. (2005) also investigated the neural response to auditory comprehension of sentences with grammatical violations using MEG. Based on Kwon’s findings, we looked for responses to grammatical violations in the 400ms and 600ms time ranges, in the left frontal and temporal lobes. The “semantic=tense” condition elicited the most significant results from controls, in the left temporal lobe from 300-500ms (Table 4c, Figure 4a) and again from 500-570ms (Table 4d, Figure 4a). This late-occurring response in the left temporal lobe occurred when comparing semantic violations to tense violations (syntactic), and may be reflective of the P600 response to syntactic violations; it may also be in response to semantic incongruences. This is consistent with Kwon’s findings, that at about 600ms, there is a response to both semantic and syntactic anomalies in the middle temporal gyrus. Though our results reveal a response occurring slightly earlier than 600ms, other reports of the P600 have found its latency to be faster in response to grammatical violations involving high-frequency words (Allen, Badecker, & Osterhout, 2003). Again, follow-up studies may want to consider the impact of lexical frequency on verb processing in agrammatism. Controls also showed sensitivity to the “semantic=tense” condition in the left frontal lobe starting at 360ms (Table 4c, Figure 4a). Kwon reports sensitivity to syntactic anomalies in inferior frontal regions at 400ms, so we may be assessing a similar response here. Finally, the “semantic=correct” condition also elicited a significant response starting at 410ms in the left temporal lobe, likely indicative of the N400 response to a semantic error. This response, to a semantic violation, is consistent with Kwon et al.’s findings (2005), as well as other reports of the N400 in response to semantic violations in spoken sentences (Mäkelä, Mäkinen, Nikkilä, Ilmoniemi, & Tiitinen, 2001).

**Table 4c:** Sentence Judgment Task Results showing pairwise contrasts, 300-500ms and 300-800ms, S=semantic violation, T=tense violation, C=correct sentence, NS=Not significant

| Controls |  |  |  | PWA Pre-Intervention |  |  | PWA Post-Intervention |  |  |
| --- | --- | --- | --- | --- | --- | --- | --- | --- | --- |
| *Left Frontal* |  |  |  |  |  |  |  |  |  |
|  | S | T | C | S | T | C | S | T | C |
| S | - | - | - | - | - | - | - | - | - |
| T | P=0.004 | - | - | NS | - | - | NS | - | - |
| C | NS | NS | - | NS | NS | - | NS | NS | - |
| *Left Temporal* |  |  |  |  |  |  |  |  |  |
|  | S | T | C | S | T | C | S | T | C |
| S | - | - | - | - | - | - | - | - | - |
| T | P=0.0004 | - | - | NS | - | - | p=0.03 | - | - |
| C | P=0.02 | NS | - | NS | NS | - | NS | NS | - |
| *Right Temporal* |  |  |  |  |  |  |  |  |  |
|  | S | T | C | S | T | C | S | T | C |
| S | - | - | - | - | - | - | - | - | - |
| T | P=0.003 | - | - | p=0.00 | - | - | NS | - | - |
| C | - | - | - | NS | NS | - | NS | NS | - |

**Figure 4a:**
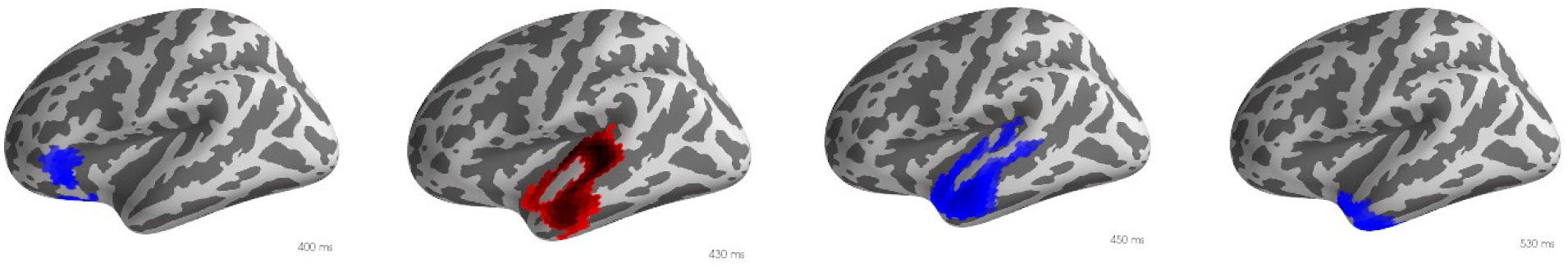
MEG response to sentence judgment task in healthy controls. a) semantic=tense, 400ms, left frontal, p=0.004, b) semantic>correct, 430ms, left temporal, p=0.02, c) semantic>tense, left temporal, 450ms, p=0.0004, d) semantic>tense, left temporal, 530ms, p=0.049.

Persons with aphasia did not show any of the same left hemisphere sensitivities to the pre-intervention grammaticality judgment task that healthy controls did. As mentioned earlier, word level processing is inherently a part of sentence processing, and if PWA were experiencing difficulty in parsing morphologically complex verbs or identifying nonwords in the lexical decision task, they would inevitably have trouble processing violations involving verb structure at the sentence level. Though agrammatism is typically described as a syntactic deficit, the idea that word level deficits are inherited into sentence processing is not new. From a psycholinguistic standpoint, it has been argued that the same processing mechanisms are responsible for the resolution of both lexical and syntactic ambiguity, and that syntactic ambiguity is actually the downstream effect of word-level lexical ambiguities (MacDonald, Pearlmutter, & Seidenberg, 1994). Studies have demonstrated that a verb’s semantic constrains influence the ease of resolution of ambiguous syntactic structures (Garnsey, Pearlmutter, Myers, & Lotocky,1997; Trueswell, Tanenhaus, & Garnsey, 1994).

That being said, though the most likely explanation is a deficit that extends to word-level processing, we are not suggesting that there is no sentence level deficit in agrammatism. It is of course possible that PWA are impaired in *both* word level processing of inflected verbs, and have difficulty in using verb inflections within a sentence context, especially considering that fact that post-intervention, the grammaticality judgment task showed differing responses, but the lexical decision task did not.

Agrammatism is a complex disorder, and a variety of factors may be contributing to sentence-level impairments. Authors have cited co-occurrence frequency (Duffield, 2016), word-form frequency (Faroqi-Shah & Thompson, 2004), frequency of verb use in discourse (Centeno & Anderson, 2011; Centeno & Obler, 2001), and the functions served by verb inflections (Friedmann & Grodzinsky, 1997; Nanousi et al., 2006; Wenzlaff and Clahsen, 2004) as factors that all might influence verb inflection processing in a sentence context. These results support the possibility that persons with aphasia have difficulty in determining the correct pairing between a verb inflection and the temporal context in a sentence (Faroqi-Shah & Dickey, 2009). This would mean that PWA have an impairment in sentence level processing of verb inflections that is separate and distinct from word level processing, and that was somewhat remediated by a morphosemantic intervention.

Finally, we also investigated the possibility of right hemisphere activation in PWA pre-intervention to determine if right hemisphere compensation was involved in verb morphology processing. A strong right temporal effect was found for PWA in the semantic=tense comparison at 310ms (Table 4c). We do not interpret this effect as right hemisphere compensation for language skills, as a nearly identical effect was seen in controls (Table 4c). However, this effect is consistent with other literature supporting right hemisphere involvement in language tasks requiring diffuse and divergent semantic processing, including processing of unusual or irrelevant semantic features for a context (Faust & Lavidor, 2003; Jung-Beeman, 2005), as in our “semantic anomaly” condition (Table 3).

### Mechanisms of Treatment-Induced Recovery

Our second research question was about the mechanisms by which persons with aphasia improved with skilled intervention addressing verb inflection deficits. All participants underwent a morphosemantic verb intervention. This intervention targeted sentence-level matching between verbs and a temporal context, and did not involve oral production. To answer this question, we looked for significant changes in neural responses to the lexical decision and sentence judgment tasks. We hypothesized that a significant change from pre- to post-intervention within one or both of the tasks would reveal the linguistic level at which the participants were improving in verb inflection processing.

Persons with aphasia did not show any changes in their neural response to the lexical decision task from pre- to post-intervention. Thus, it was concluded that documented behavioral improvements post-intervention were not occurring via word-level processing. However, PWA did begin to show significant responses to the sentence judgment task after intervention (Figure 4b). They showed significant responses to the “semantic=tense” condition at 720ms in the left temporal lobe (Table 4d), and a significant response to the same condition at 960ms in the right temporal lobe. Persons with aphasia did not show sensitivity to any of the sentence task conditions in the frontal lobe at any time point.

**Figure 4b:**
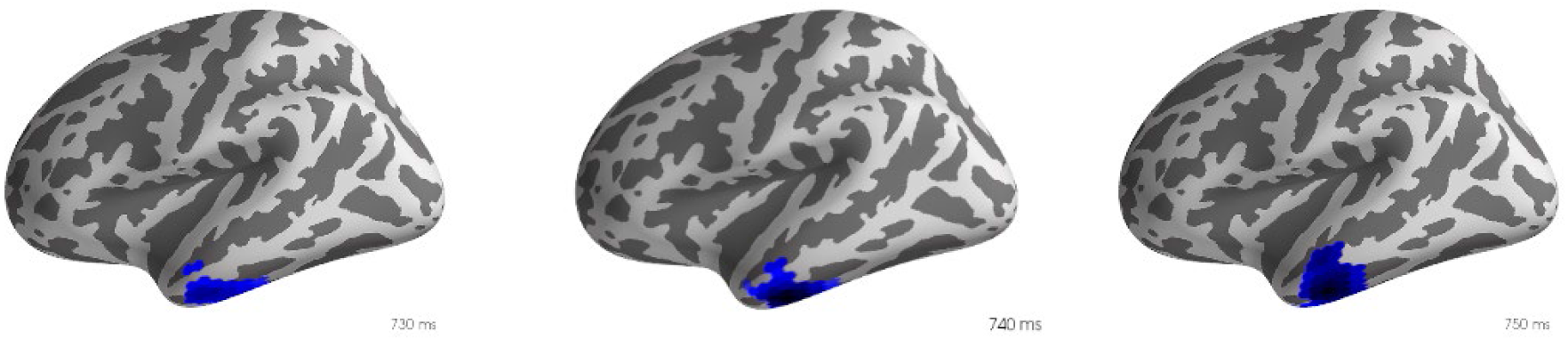
MEG response to sentence judgment tasks in PWA post-intervention. All figures capture significant clusters in response to the semantic>tense comparison in the left temporal lobe (p=0.03). From left to right, 730, 740, and 750ms.

**Table 4d:** Sentence Judgment Task Results showing pairwise contrasts, 500-700 and 500-1000ms

| Table 1a: Sentence Judgment Task Results showing pair-wise contrasts, 500-700 and 500-1000ms |  |  |  |  |  |  |  |  |  |
| --- | --- | --- | --- | --- | --- | --- | --- | --- | --- |
| Controls |  |  | PWA Pre-Intervention |  |  | PWA Post-Intervention |  |  |  |
| *Left Frontal, 500-700ms* |  |  |  |  |  |  |  |  |  |
|  | S | T | C | S | T | C | S | T | C |
| S | - | - | - | - | - | - | - | - | - |
| T | NS | - | - | NS | - | - | NS | - | - |
| C | NS | NS | - | NS | NS | - | NS | NS | - |
| *Left Temporal, 500-700ms* |  |  |  |  |  |  |  |  |  |
|  | S | T | C | S | T | C | S | T | C |
| S | - | - | - | - | - | - | - | - | - |
| T | p=0.049 | - | - | NS | - | - | NS | - | - |
| C | NS | NS | - | NS | NS | - | NS | NS | - |
| *Right temporal, 500-1000ms* |  |  |  |  |  |  |  |  |  |
|  | S | T | C | S | T | C | S | T | C |
| S | - | - | - | - | - | - | - | - | - |
| T | p=0.026 | - | - | NS | - | - | p=0.031 | - | - |
| C | NS | NS | - | NS | NS | - | NS | NS | - |

A lack of a response in the frontal lobe is not unexpected, given most of participants (and most persons with Broca’s aphasia) have left frontal lesions. Control data in our experiment indicates that the left frontal and left temporal lobes both play crucial roles in processing during the sentence judgment tasks, and that, like healthy controls, PWA recruit left temporal regions for processing grammatical violations in sentences.

It was expected that morphosemantic intervention for agrammatism would operate via improving participants’ sentence-level processing of verbs, given that the intervention involved sentence level comprehension tasks. This also fits with findings indicating that comprehension of verb inflections within a given syntactic context is impaired (Faroqi-Shah & Dickey, 2009), however, interestingly, all participants (PWA and healthy controls) only showed significant responses to the morphosemantic (“tense”) violation condition when it was statistically compared to the semantic condition. So, the brain responses to different categories of violations were significantly different, though the responses to individual violations (with the exception of semantic violations in controls) did not differ from the responses to correct sentences. However, we can conclude from this finding that for both healthy speakers and PWA, there is indeed something unique about “morphosemantic” (tense) violations (Faroqi-Shah & Dickey, 2009), that is not the same as general “wrongness” of a sentence.

Though the neural responses to the semantic=tense test condition were similar in PWA and healthy controls, PWA’s response was about 200ms delayed compared to healthy controls. This is consistent with findings of delayed responses in PWA to single word stimuli (Zipse et al., 2011) and delayed left temporal responses during receptive language tasks (Breier, 2004). Though this is the first study we know of to reveal a delay in neural responses to sentence level auditory comprehension tasks in PWA, single word processing delays would inherently influence sentence comprehension as well.

We also investigated right hemisphere processing in PWA post-intervention, to account for the possibility of right hemisphere compensation associated with treatment, as has been reported in other accounts of aphasia recovery (Breier et al., 2011; Schlaug, Marchina, & Norton, 2009). We did find a late (960ms) right temporal response post-intervention, but not pre-intervention. However, we then confirmed that the same response (940ms) was actually occurring in healthy controls. The late nature of this effect implies that it may not be directly related to the task judgment, and rather may reflect a non-syntactic “sentence wrap-up” effect, an effect that has been robustly documented in reading paradigms using eye tracking in healthy controls (Hirotani, Frazier, & Rayner, 2006; Warren, White, & Reichle, 2009), and auditory comprehension tasks (Hagoort, 2003), and also noted in persons with aphasia (Balogh, Zurif, Prather, Swinney, & Finkel, 1998). Furthermore, the pre-intervention right temporal effect we identified in PWA (Table 4c) was not present after treatment. This is actually consistent with reports of decreased right hemisphere activation and increased perilesional activation associated with stronger recovery with therapy (Breier et al., 2010).

Overall, it seems that PWA’s post-intervention sentence level processing mirrors that of healthy controls (Figure 2), and improvements are occurring via continued activation of pre-morbid language processing centers, rather than right hemisphere compensatory activity. This is consistent with a recent review of neuroimaging literature, which reported that there is minimal evidence supporting the role of non-language areas in post-stroke language recovery (Harwigsen, 2017).

There is one final aspect to our findings that must be resolved: though PWA showed impaired language processing during the lexical decision (word-level) task, improvement post-intervention was associated with changes in processing during the grammaticality judgment (sentence-level) task. This supports the possibility of a sentence-level deficit that not entirely linked to the word-level. This would mean that PWA experience deficits in verb inflection processing at *both* the word and sentence level, and our intervention only targeted one level (sentence). As stated earlier, a morphosemantic impairment has been well documented in this population (Faorqi-Shah & Dickey, 2009), and a word level verb inflection processing deficit is not mutually exclusive with a morphosemantic impairment.

### Conclusions

Overall, the findings of this experiment are unique in that they posit the possibility that deficits in agrammatism exist at the single word level and may not be purely syntactic in nature. These findings should be interpreted with caution, as only 6 people participated in this study, and we also found impaired processing pre-intervention in PWA at the sentence level.

Despite finding evidence for a word level deficit, this study also provides support for the efficacy of sentence-level training programs for agrammatism. Our findings provide further support for the efficacy of a morphosemantic verb therapy for verb morphology deficits (Faroqi-Shah, 2008), and indirectly supports the value of other sentence level interventions (Thompson, et al., 2010).

Limitations of this study include a relatively small sample size for PWA, and the lack of a true age-matched control for PWA. Though data was originally collected from an age-matched sample, external noise in the MEG recordings prevented us from assessing a true signal from their brain scans, and comparisons to neurologically adults relied heavily on the signal collected from young adults. We are aware that aging effects in the PWA may have confounded the results we assessed, given some literature has pointed to the possibility that syntactic processing specifically declines with age (Obler, Fein, Nicholas, & Albert, 1991). However, a more recent ERP study investigating the neural response to syntactic violation in aging adults found no difference in the amplitude or latency of the P600 in older adults, though it did report slight differences in spatial distribution (Kemmer, Coulson, De Ochoa, & Kutas, 2004). Additionally, recent literature has indicated that word-level comprehension remains relatively stable in healthy aging (Abrams & Farrell, 2011). Of course, future research should aim to replicate our findings with a more closely age-matched control group to account for potential effects of aging.

## Acknowledgements

Part of the analysis reported in this paper was completed as part of the first author’s Masters Thesis.

## Appendix 1 Participant Language Profiles

**Table 5:**
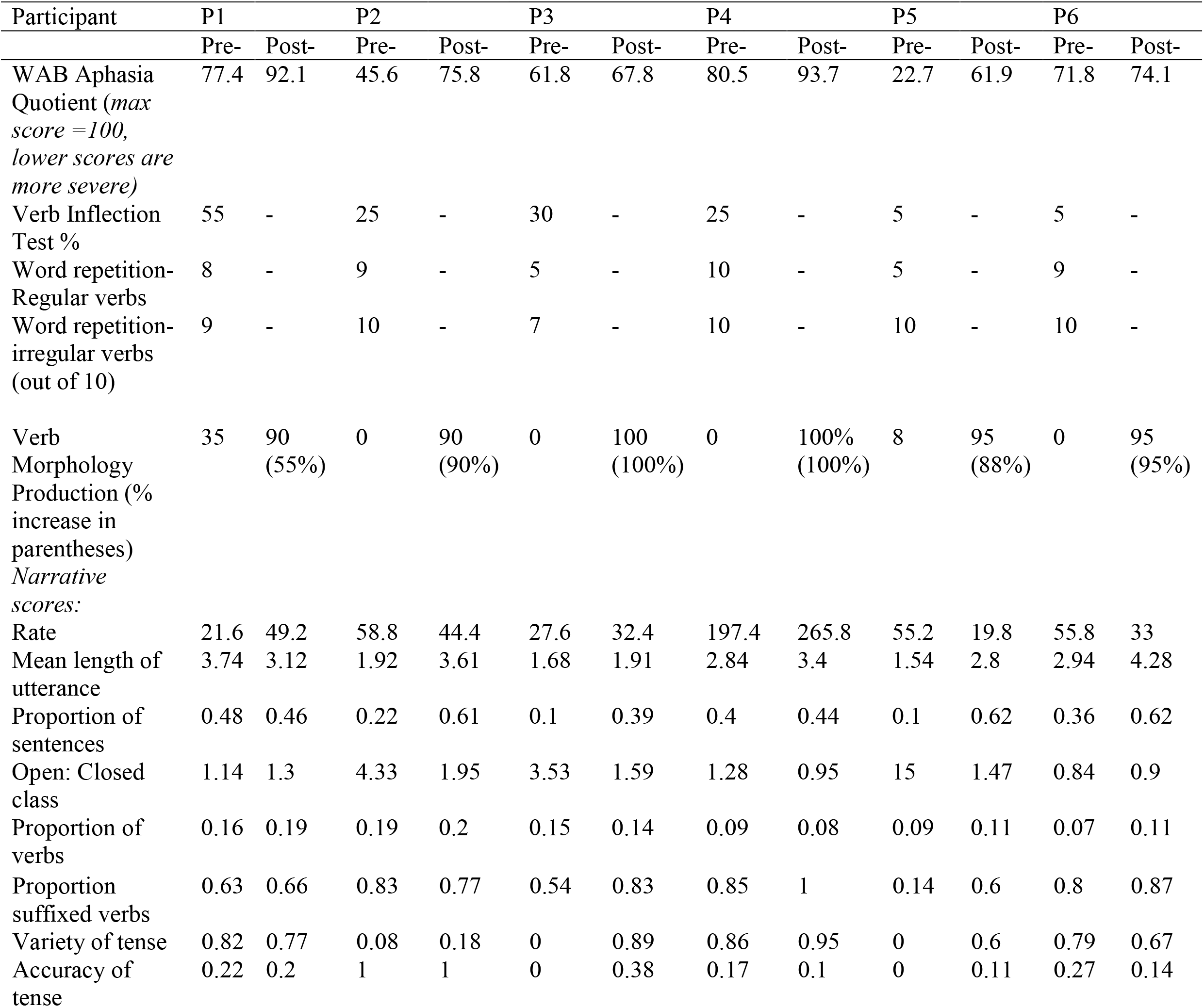
Participant language profiles

## Notes

### Competing Interest Statement

The authors have declared no competing interest.

